# Hydroxyurea inhibits ERAD-L independently of S-phase arrest in budding yeast

**DOI:** 10.1101/2023.01.22.525111

**Authors:** Yuki Takano, Kunio Nakatsukasa

## Abstract

Misfolded luminal and membrane proteins in the endoplasmic reticulum (ER) are recognized and retrotranslocated to the cytosol for proteasomal degradation, a process referred to as ER-associated degradation (ERAD). In *Saccharomyces cerevisiae*, ERAD substrates with luminal lesions are targeted for proteasomal degradation by the Hrd1 ubiquitin ligase complex (ERAD-L pathway). Membrane proteins containing lesions within their membrane-spanning regions are also targeted for degradation by the Hrd1 complex (ERAD-M pathway), while those containing lesions within their cytosolic regions are targeted for degradation mainly by the Doa10 ubiquitin ligase complex (ERAD-C pathway). Here, we demonstrate that hydroxyurea (HU), which is widely used to arrest cells in S-phase and is also used to manage several diseases including sickle cell anemia and chronic myeloproliferative disorders, inhibited ERAD-L, but not ERAD-M or -C. HU-mediated inhibition of ERAD-L occurred independently of S-phase arrest. In HU-treated cells, the integrity of the Hrd1 ubiquitin ligase complex remained intact and substrate recognition was unaffected. Moreover, induction of the unfolded protein response was undetectable in cells in which ERAD-L was inhibited by HU. These results suggest an unexpected action of HU, which modulates protein quality control in the secretory pathway, and also suggest the existence of an additional regulatory step in ERAD.

## INTRODUCTION

Misfolded proteins accumulated in the endoplasmic reticulum (ER) elicit the unfolded protein response (UPR), which activates an expansive gene expression program to restore ER homeostasis (1,2). Genes induced by the UPR include those encoding the components of ER-associated degradation (ERAD) machinery, by which misfolded proteins in the ER are recognized, retrotranslocated to the cytosol, ubiquitinated, and degraded by the proteasome (3–13). In budding yeast, ERAD substrates with luminal lesions are targeted to the Hrd1 ubiquitin ligase complex (ERAD-L pathway). Membrane proteins containing lesions within their membrane-spanning regions are also targeted to the Hrd1 ubiquitin ligase (ERAD-M pathway). By contrast, membrane proteins with misfolded lesions facing the cytosol are primarily recognized by Doa10, another integral membrane ubiquitin ligase in the ER membrane (14–16). Ubiquitinated substrates are then segregated from the ER membrane and delivered to the proteasome for degradation by the action of the Cdc48/p97 AAA-ATPase (4,17–19).

Given the critical role of ERAD in regulation of ER homeostasis and its relevance to numerous diseases (20), it is conceivable that defects in this process can significantly impact cell viability. Indeed, the ERAD pathway is a potential target for pharmacological intervention with tumors (20–22). Several small molecules that inhibit ERAD have been reported. For example, inhibition of ER-mannosidase I stabilizes some misfolded glycoproteins (23). Manipulation of the redox potential by addition of alkylating reagents, such as *N*-ethylmaleimide and diamide, blocks the retrotranslocation of several misfolded proteins (24). Eeyarestatin 1 inhibits ERAD by targeting cytosolic p97 (25–27), although it also impairs Sec61-dependent protein translocation into the ER, vesicular transport within the endomembrane system, and Ca^2+^ homeostasis (28–30).

Since it was first synthesized more than 150 years ago, HU has been widely used in both basic research and clinical practice (31,32). HU has been a useful agent for studying the cell cycle due to its ability to inhibit DNA replication by inactivating ribonucleotide reductase (RNR), and consequently to induce S-phase arrest and checkpoint activation (33–35). In clinical practice, HU was first reported to have antitumor activity in the 1960s and has become established as a drug used to treat a variety of diseases, including brain tumors (36), chronic myeloproliferative disorders (37), and sickle cell anemia (38–40). Recently, HU has been shown to improve spatial memory in a mouse model of Alzheimer’s disease, making it a promising drug to treat cognitive decline in this disease (41). However, the mechanism by which HU inhibits RNR is poorly understood (32,42). In addition, HU affects other enzymes due to its less specific action, which may further affect cellular responses by unidentified factors. For example, HU appears to target iron-sulfur (Fe-S) centers *in vivo* by producing reactive oxygen species (43). HU is also degraded over time and in the presence of heat to generate *N*-hydroxyurethane, hydrogen cyanide, nitric oxide, and peroxides (32,37,44–46). Thus, despite the evidence that HU is an effective agent against a variety of diseases, its mechanisms of action, side effects, and toxicity remain largely unclear. It is important to further elucidate the effects of HU on cellular function to use it as a reliable agent for treatment of both oncologic and nontumor diseases.

In the course of analyzing the possible regulation of ERAD during the cell cycle, we found that HU inhibits ERAD-L, but not ERAD-M or -C, independently of S-phase arrest. These results suggest an unexpected action of HU and also suggest the existence of an unidentified regulatory step in ERAD.

## RESULTS and DISCUSSION

### HU inhibits ERAD-L

We initially sought to examine whether ERAD is physiologically regulated during the cell cycle. To this end, we performed cycloheximide chase analysis of model ERAD substrates to monitor their degradation in cells synchronized at G1, S, or G2/M phase by treatment with α-factor, HU, or nocodazole (NC), respectively. The cell cycle synchronization was confirmed by the levels of Sic1 and Clb2 (Fig. S1A), which peak at G1 and G2/M phases, respectively (47). Model ERAD-L substrates including CPY*, a soluble substrate due to the presence of a missense mutation in an otherwise vacuole-targeted protease (48), KHN, a heterologously expressed simian virus 5 hemagglutinin neuraminidase (HN) that is fused with the cleavable signal sequence from the yeast Kar2 (the ER luminal Hsp70) (16), and KWW, a chimeric protein comprising KHN luminal domain/Wsc1 transmembrane domain/Wsc1 cytosolic domain (16), (Fig. 1A) were degraded in cells treated with α-factor or NC to a similar extent as in control cells (Fig. 1B–D, lanes 5–8 and 13–16). However, their degradation was strongly inhibited in cells treated with HU (Fig. 1B–D, lanes 9–12). By contrast, the turnover of model ERAD-M substrates including 6myc-Hmg2, the yeast HMG-CoA reductase isozyme (49), and Pdr5*, a 12 transmembrane protein that harbors misfolded lesions near these domains (50,51), (Fig. 2A) was unaffected by all of these drugs (Fig. 2B–C, Fig. S1B). 3HA-tagged Pdr5* instead of the originally used 1HA-tagged Pdr5* (50,51) was used to better detect Pdr5*. Degradation of 3HA-Pdr5* was dependent on Hrd1 but independent of Der1 (Fig. S2), as suggested previously for 1HA-Pdr5* (14,51). Finally, the turnover of model ERAD-C substrates including Ste6*, a C-terminal truncated version of the a-factor transporter Ste6 (52) and Pca1, a cadmium transporting P-type ATPase whose proteasome-dependent degradation is exclusively dependent on Doa10 (53), (Fig. 2A) was also unaffected by all of these drugs (Fig. 2D–E, Fig. S1B). These results demonstrate that only ERAD-L, not ERAD-M or -C, is inhibited by administration of HU.

**Fig. 1:**
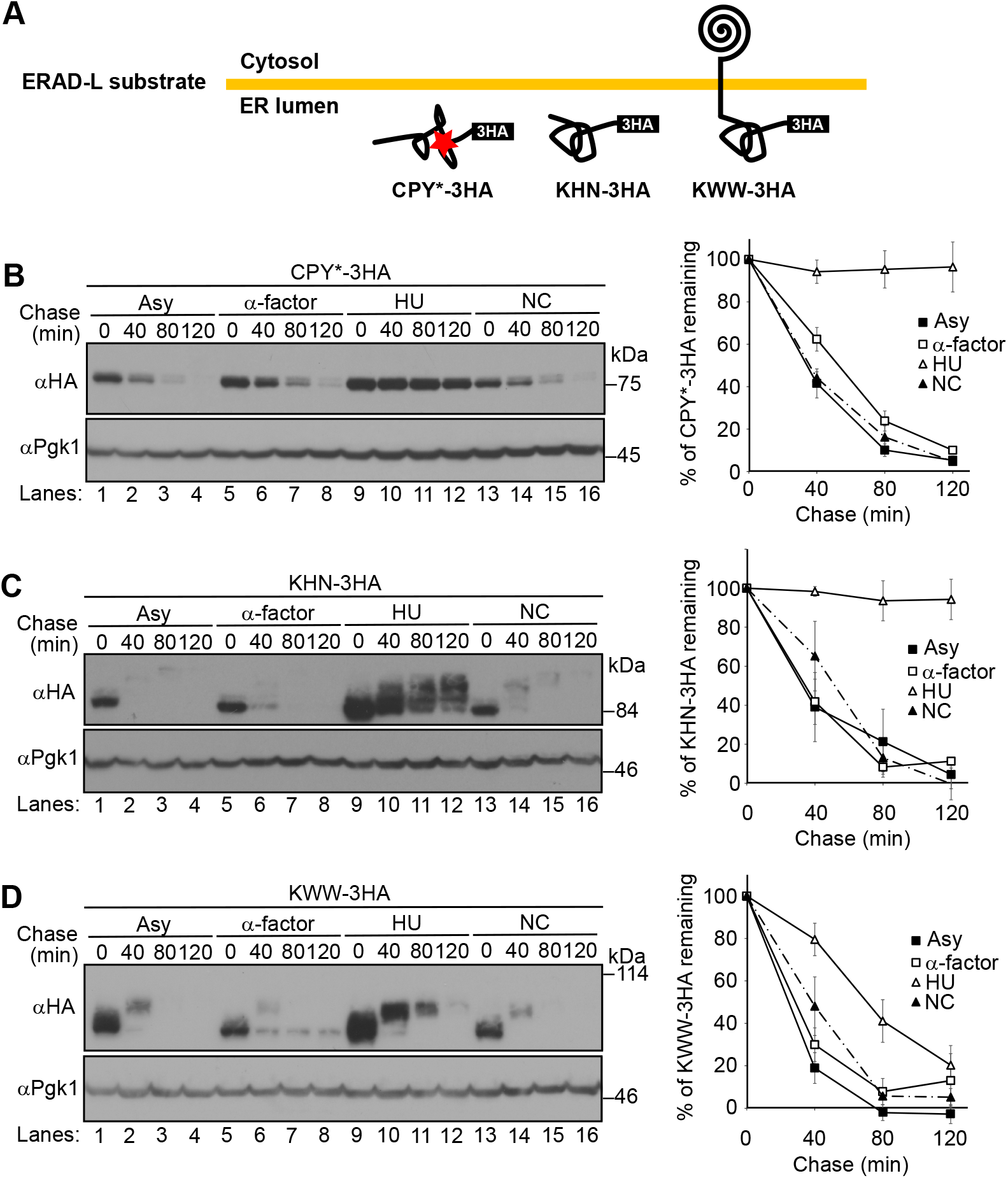
Cycloheximide chase analysis of ERAD-L substrates in cells synchronized at G1, S, or G2/M phase. **(A)** Schematic representation of model ERAD-L substrates. **(B)** WT cells expressing CPY*-3HA were grown at 30°C and synchronized at G1, S, or G2/M phase by incubation with α-factor, HU, or NC, respectively, for 2.5 hr. Cycloheximide (200 μg/mL) was added and cells were collected at the indicated time points. Total cell lysates were subjected to western blotting with an anti-HA antibody. Pgk1 served as a loading control. CPY*-3HA signals were normalized to Pgk1 signals. Quantification of the results is shown. The data represent the mean ± SE of three independent experiments. **(C–D)** Cycloheximide chase analyses of KHN-3HA (C) and KWW-3HA (D) were performed as in (B). Signals for each substrate were normalized to Pgk1 signals. Quantification of the results is shown. The data represent the mean ± SE of three independent experiments.

**Fig. 2:**
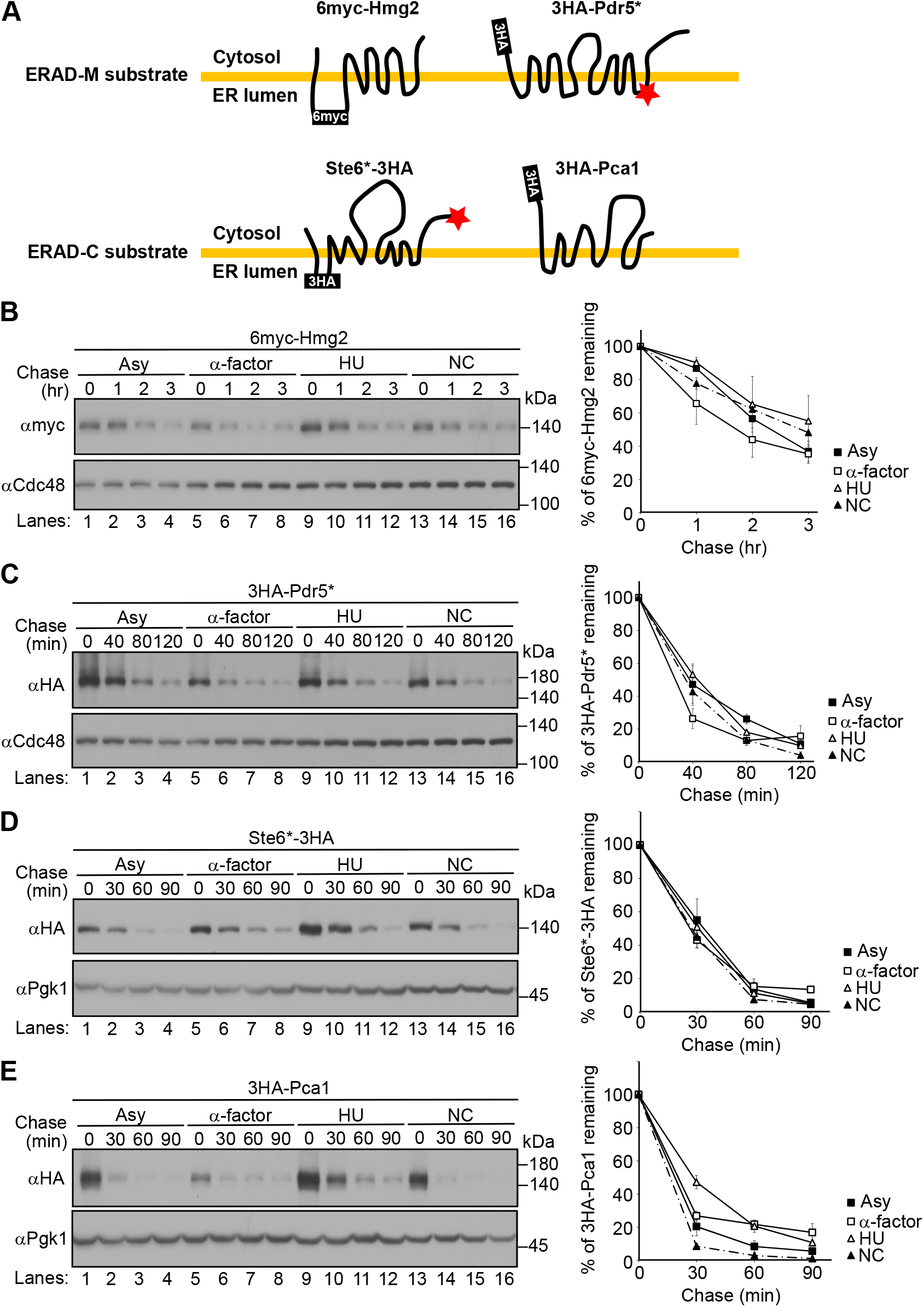
Cycloheximide chase analysis of ERAD-M and -C substrates in cells arrested at G1, S, or G2/M phase. **(A)** Schematic representation of model ERAD-M and -C substrates. **(B–E)** Cycloheximide chase analyses of 6myc-Hmg2 (B), 3HA-Pdr5* (C), Ste6*-3HA (D), and 3HA-Pca1 (E) in cells arrested at G1, S, or G2/M phase were performed as in Fig. 1B. Expression of Ste6*-3HA was induced under the control of the *GAL1* promoter in medium containing 2% galactose as a sole carbon source. Cdc48 (B–C) and Pgk1 (D– E) served as loading controls. Quantification of the results is shown. Signals for each substrate were normalized to Cdc48 or Pgk1 signals. The data represent the mean ± SE of three independent experiments.

### HU inhibits ERAD-L independently of S-phase arrest

To investigate whether S-phase synchronization is a prerequisite for HU-mediated inhibition of ERAD-L, we analyzed CPY* degradation under several conditions. First, we found that CPY* degradation was strongly inhibited even in cells treated with HU only for 15 min (Fig. 3A, lanes 17–20). This is much shorter than ~2.5 hr, which is generally needed to fully synchronize the cell cycle of budding yeast. Second, we analyzed the level of CPY* over time during the cell cycle. Cells were synchronized at G1 phase by treatment with α-factor and then the cell cycle was restarted by removing α-factor. Cell cycle progression was confirmed by the levels of Sic1 and Clb2 (Fig. 3B, αSic1 and αClb2). The level of CPY* was unchanged during the cell cycle (Fig. 3B, αHA). Third, under our experimental conditions, cells entered S-phase at 30 min after the cell cycle was restarted from G1 phase (Fig. 3B, lane 4, indicated by arrow; Fig. 3C, upper panel, lane 6). In these cells, CPY* was robustly degraded (Fig. 3C, lower panel, αHA, lanes 9–12). Fourth, the DNA-damaging alkylating agent methyl methanesulfonate (MMS), which also induces genotoxic stress and activates the S-phase checkpoint, did not stabilize CPY* (Fig. 3D). Together, these results suggest that HU inhibits ERAD-L independently of S-phase arrest.

**Fig. 3:**
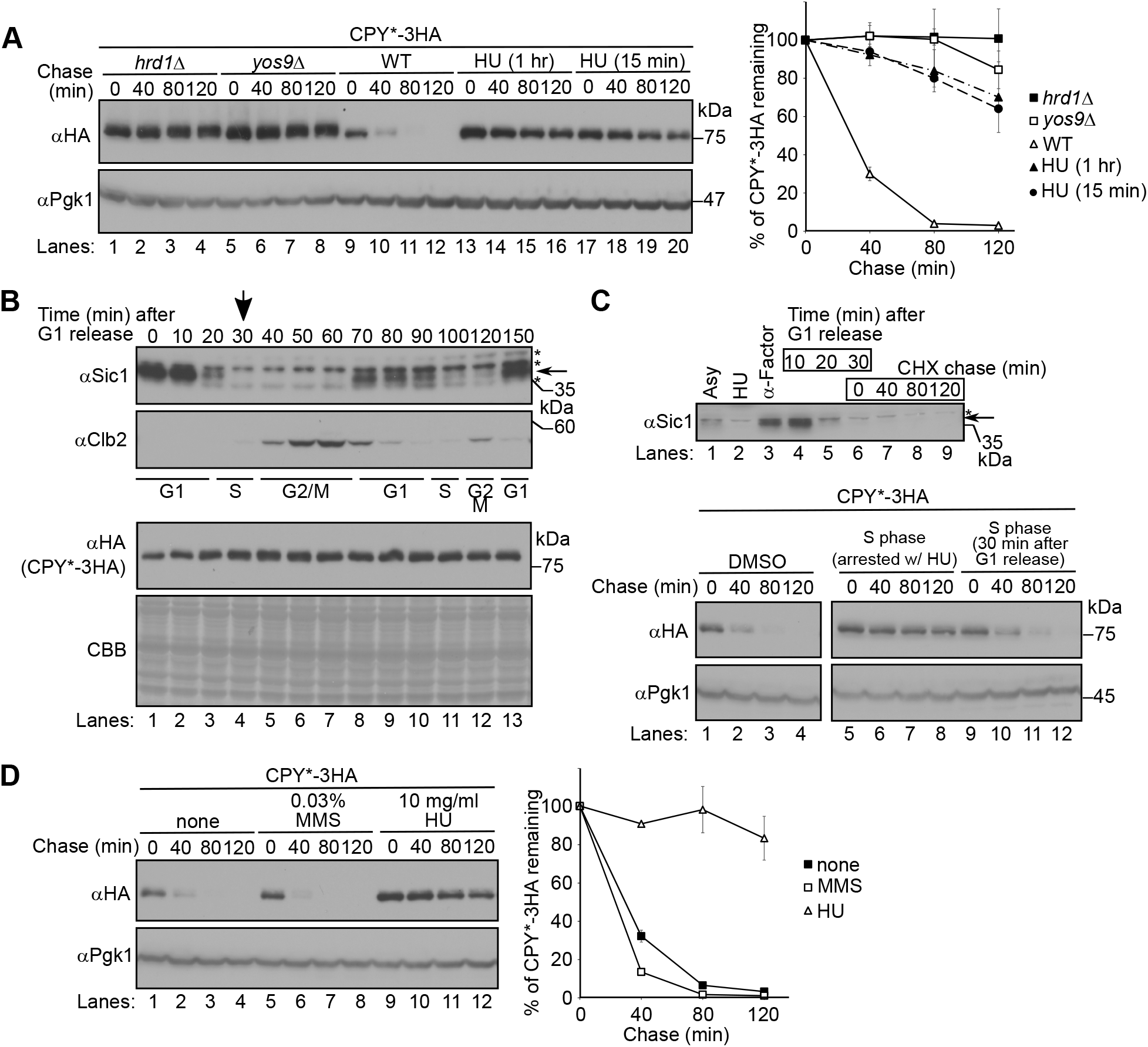
HU inhibits ERAD-L independently of S-phase arrest. **(A)** Cycloheximide chase analysis of CPY*-3HA was performed as in Fig. 1B. Where indicated, cells were treated with 10 mg/mL HU for 1 hr (lanes 13–16) or 15 min (lanes 17–20) before the addition of cycloheximide. CPY*-3HA signals were normalized to Pgk1 signals. Quantification of the results is shown. The data represent the mean ± SE of three independent experiments. **(B)** Cells expressing CPY*-3HA were grown at 30°C until OD_600_ reached ~0.35 before being synchronized at G1 phase with α-factor. After 2.5 hr, α-factor was removed and the cell cycle was restarted in fresh medium (0 min). At the indicated time points, an equal amount of cells was collected by centrifugation and subjected to western blotting with an anti-Sic1, anti-Clb2, or anti-HA (CPY*) antibody. Coomassie Brilliant Blue (CBB) staining of the membrane served as a loading control. Cells that entered S-phase (indicated by the arrow) were used for the cycloheximide chase analysis in (C). **(C)** Cells expressing CPY*-3HA were grown to log phase (upper panel, lane 1) and arrested at G1 phase (lane 3), and the cell cycle was restarted as in (B). After 30 min (lane 6), when cells entered S-phase, cycloheximide chase analysis of CPY* was performed as in Fig. 1B (lower panel, lanes 9–12). Cell cycle arrest and progression were monitored by the level of Sic1 (upper panel). Cycloheximide chase analysis of CPY* in cells arrested at S-phase by treatment with HU for 2.5 hr was also performed (lower panel, lanes 5–8). **(D)** Cycloheximide chase analysis of CPY*-3HA was performed as in Fig. 1B. Where indicated, cells were treated 0.03% MMS or 10 mg/mL HU for 1 hr. CPY*-3HA signals were normalized to Pgk1 signals. Quantification of the results is shown. The data represent the mean ± SE of three independent experiments.

### The integrity of the Hrd1 ubiquitin ligase complex and the recognition of ERAD-L substrates are unaffected in HU-treated cells

To investigate the mechanism by which HU inhibits ERAD-L, we investigated if treatment with HU alters the integrity of the Hrd1 complex (Fig. 4A). The functional tagged form of Hrd1-3FLAG (54) was immunoprecipitated from digitonin-solubilized membranes prepared from cells treated with HU. The components of the Hrd1 complex including Usa1, Hrd3, and Der1 as well as Yos9 (55–58) were all co-precipitated to the same extent from HU-treated and untreated cells (Fig. 4B). Der1 was expressed slightly more in HU-treated cells than in non-treated cells (Fig. 4B, compare lanes 2 and 3, αDer1), but previous studies suggested that overexpression of Der1 does not significantly interfere with the degradation of luminal substrates (59–62). Therefore, induction of Der1, if any, did not explain HU-mediated inhibition of ERAD-L. These results suggest that HU does not affect the integrity of the Hrd1 complex.

**Fig. 4:**
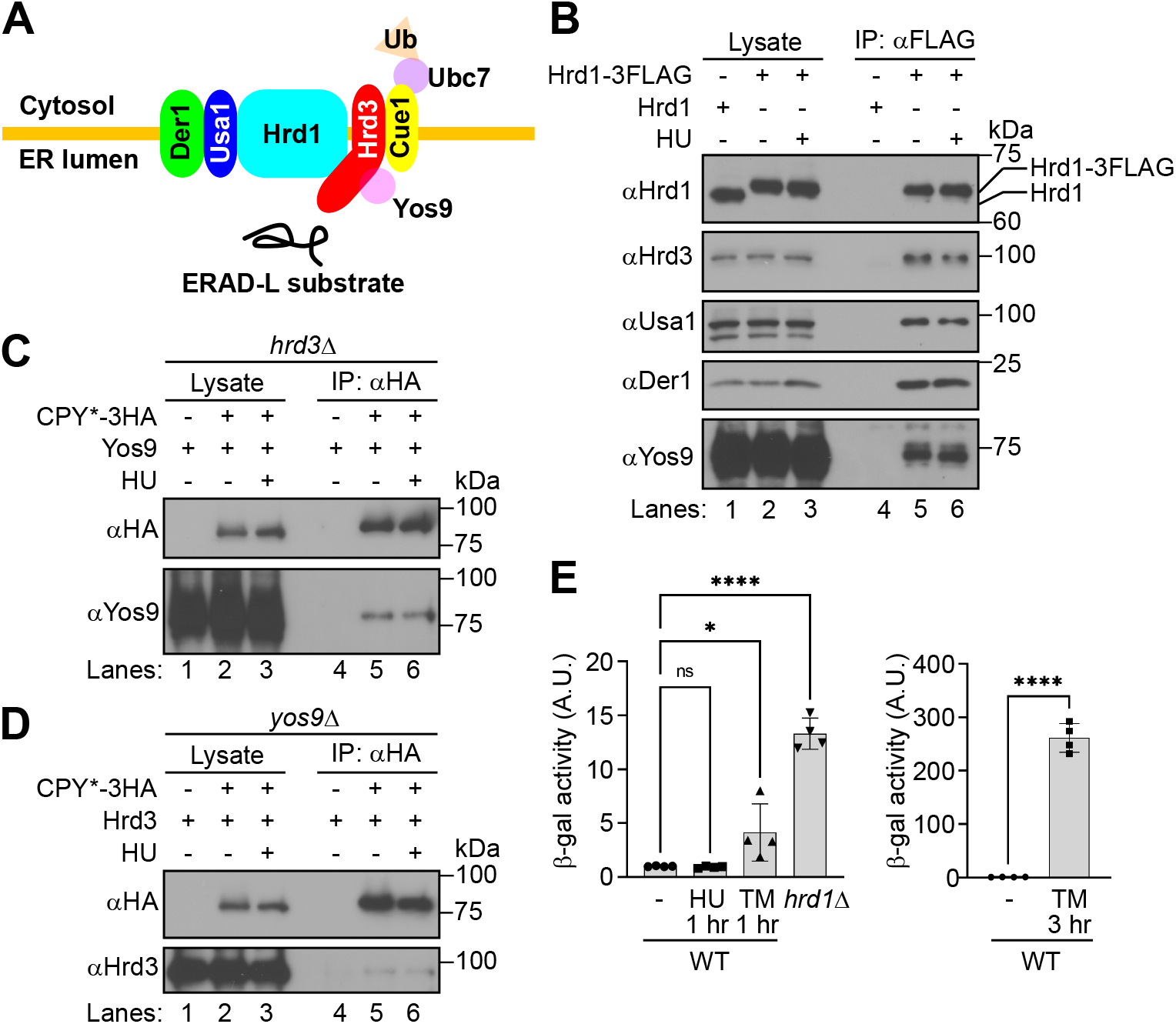
HU does not affect substrate recognition or the integrity of the Hrd1 complex. **(A)** Schematic representation of the Hrd1 complex. **(B)** Hrd1-3FLAG was immunoprecipitated from cells treated with (+) or without (-) 10 mg/mL HU for 1 hr at 30°C. The immunoprecipitated materials were analyzed by western blotting with the indicated antibodies. **(C–D)** CPY*-3HA was immunoprecipitated with an anti-HA antibody from *hrd3Δ* cells (C) or *yos9*Δ cells (D) treated with (+) or without (-) 10 mg/mL HU for 1 hr at 30°C. The immunoprecipitated materials were analyzed by western blotting with the indicated antibodies. **(E)** The extent of UPR induction was measured by the β-galactosidase assay as previously described (73,74). Where indicated, cells were treated with 10 mg/mL HU for 1 hr or 10 μg/mL tunicamycin for 1 or 3 hr. The data represent the mean ± SD of four measurements (two experiments using independent overnight cultures performed in duplicate); ns, not significant; *p < 0.05; ****p < 0.0001; one-way ANOVA with Dunnett’s multiple comparison test (left panel); unpaired t-test (right panel).

We next analyzed substrate recognition in cells treated with HU. Yos9 and Hrd3 recognize ERAD-L substrates independently of each other (56,63). When CPY* was immunoprecipitated from the digitonin-solubilized membrane fraction prepared from HU-treated *hrd3Δ* cells, Yos9 was co-immunoprecipitated to the same extent as from non-treated *hrd3Δ* cells (Fig. 4C). Similarly, Hrd3 was co-immunoprecipitated with CPY* to the same extent from HU-treated and non-treated *yos9Δ* cells (Fig. 4D). These results suggest that HU does not affect Yos9/Hrd3-mediated recognition of ERAD-L substrates.

Pdr5* harbors two *N*-linked glycans and its degradation is dependent on Yos9 (64) (Fig. S3). However, HU did not inhibit Pdr5* degradation (Fig. 2C), again supporting the idea that HU does not affect Yos9-mediated substrate recognition. In addition, ERAD-L substrates may be recycled between the ER and Golgi before their degradation. Blockade of ER-Golgi transport reportedly inhibits ERAD-L *in vivo* (16,65,66). However, in this study, when degradation of KHN and KWW was inhibited by HU administration, they acquired *O*-linked glycosylation, which was evident from their slowed migration in the blots, due to their persistent recycling between the ER and Golgi during the chase period (16,66) (Fig. 1C and D). This result suggests that these substrates were recycled persistently between the ER and Golgi when their degradation was inhibited by HU. Thus, HU-mediated inhibition of ERAD-L does not appear to be due to blockade of ER-Golgi transport.

### The UPR is hardly induced in HU-treated cells

A previous study suggested that strong induction of the UPR by treatment of cells with the ER-specific stressor tunicamycin, which blocks *N*-linked glycosylation, or the reducing agent DTT inhibits ERAD-L (67). We therefore investigated if HU inhibits ERAD-L by inducing the UPR. However, while the UPR was induced in cells treated with tunicamycin, treatment with HU at the concentration that was sufficient to block ERAD-L hardly activated the UPR (Fig. 4E, left panel). The extent of UPR induction was lower in HU-treated cells than in *hrd1Δ* cells, in which the UPR is chronically induced due to the absence of ERAD-L (68,69). The level of UPR induction was only ~4-fold higher in cells treated with tunicamycin for 1 hr than in untreated cells, but the UPR was significantly induced in cells treated with tunicamycin for 3 hr (right panel). These results suggest that UPR induction does not explain the stabilization of ERAD-L substrates upon HU treatment.

In sum, we found that HU inhibits ERAD-L but not ERAD-M or -C. How does HU specifically inhibit ERAD-L? In the latter half of the ERAD pathway, ERAD-L, -M, and -C all rely on the actions of Cdc48/p97 and proteasomes. However, ERAD-M and -C were largely unaffected by HU (Fig. 2). Thus, HU inhibits ERAD-L probably by compromising the steps in the first half of the ERAD pathway, which is specific to ERAD-L. Importantly, the recognition of ERAD-L substrates by Yos9/Hrd3 in the ER lumen and the integrity of the membrane-associated Hrd1 complex were unaffected (Fig. 4). One attractive hypothesis is that HU directly or indirectly inhibits the movement of ERAD-L substrates from the ER lumen to the cytosol through the Hrd1 complex. It is also possible that the ubiquitination state of ERAD-L substrates is quantitatively and/or qualitatively changed in the presence of HU. Clearly, the detailed mechanism underlying HU-mediated ERAD-L inhibition needs to be further investigated in the future. Nonetheless, our results suggest an unexpected action of HU, which modulates protein quality control in the secretory pathway, and also suggest the existence of an additional regulatory step in ERAD.

## EXPERIMENTAL PROCEDURES

### Yeast strains, plasmids, oligonucleotide primers, and antibodies

Yeast strains, plasmids, oligonucleotide primers, and antibodies used in this study are listed in Supplementary Tables I, II, III, and IV, respectively. Cells were grown in YP-rich medium (1% yeast extract and 1% peptone) supplemented with 2% glucose (YPD) or synthetic complete medium (0.67% yeast nitrogen base without amino acids) supplemented with all standard amino acids and 2% glucose (SD). Where indicated, 2% galactose was used instead of glucose.

### Cycloheximide chase analysis and western blotting

Cycloheximide chase experiments and western blotting were performed essentially as described previously (70–72).

### UPR assay

Cells harboring pJC104 encoding 4×UPRE (four copies of the UPR element)-*lacZ* (73) were grown overnight until OD_600_ reached 0.3–0.4 in SD medium. The ß-galactosidase assay was performed as described previously (74).

### Immunoprecipitation

Co-immunoprecipitation was performed essentially as described previously (54). Briefly, cells were grown to mid-log phase (OD_600_≈1.5). Cells (~100 OD_600_) were harvested and disrupted with glass beads in lysis buffer (20 mM HEPES-KOH pH 7.4, 50 mM KOAc, 2 mM EDTA, and 0.1 M sorbitol) supplemented with a complete protease inhibitor cocktail (PIC) (Roche) by agitation on a vortex mixer. The homogenate was collected and pooled, and the beads were rinsed with buffer 88 (20 mM HEPES pH 7.4, 150 mM KOAc, 250 mM sorbitol, and 5 mM MgOAc). After unbroken cells were removed by centrifugation at 300 × g for 5 min at 4°C, the supernatant was centrifuged at 20,000 × g for 20 min at 4°C. The pellet (membrane) fractions were solubilized in solubilization buffer (20 mM Tris-HCl pH 7.5, 100 mM NaCl, and 10% glycerol) supplemented with 0.5% TritonX-100. The cleared lysate was added to anti-FLAG M2 affinity gel (Sigma-Aldrich). After nutation at 4°C for 1 hr, the gel was washed three times with solubilization buffer supplemented with 0.5% TritonX-100 before proteins were eluted with SDS-PAGE sample buffer.

## Data availability statement

No datasets were generated during the current study.

## Supporting information

This article contains supporting information.

## Acknowledgments

We thank the members of the Nakatsukasa laboratory for discussions. We also thank the National BioResource Project, Jeffrey L. Brodsky, Randolph Y. Hampton, Ernst Jarosch, Takumi Kamura, Jaekwon Lee, Susan Michaelis, Davis T.W. Ng, Thomas Sommer, Peter Walter, and Jonathan S. Weissman for plasmids, strains, and antibodies. We acknowledge the assistance of the Research Equipment Sharing Center at Nagoya City University.

## Funding

This work was supported by grants from the Toray Science Foundation to K.N., the Toyoaki Scholarship Foundation to K.N., and JSPS KAKENHI to K.N. (Grant Numbers 18K19306 and 19H02923).

## Conflict of interest

The authors declare that they have no conflicts of interest with the contents of this article.

## Notes

### Competing Interest Statement

The authors have declared no competing interest.

