## Supplemental Tables 1-4 for "Hydroxyurea inhibits ERAD-L independently of S-phase arrest in budding yeast"

**Supplementary Table I**

Yeast strains used in this study

| **Name** | **Description** | **Source** |
|---|---|---|
| BY4741 | *MAT***a** *his3Δ1 leu2Δ0 met15Δ0 ura3Δ0* | Dr. S. Michaelis |
| E3Δhrd1 | *MAT***a** *his3Δ1 leu2Δ0 met15Δ0 ura3Δ0 hrd1Δ::KanR* | Open Biosystems |
| E3Δhrd3 | *MAT***a** *his3Δ1 leu2Δ0 met15Δ0 ura3Δ0 hrd3Δ::KanR* | Open Biosystems |
| E3Δyos9 | *MAT***a** *his3Δ1 leu2Δ0 met15Δ0 ura3Δ0 yos9Δ::KanR* | Open Biosystems |
| E3Δder1 | *MAT***a** *his3Δ1 leu2Δ0 met15Δ0 ura3Δ0 der1Δ::KanR* | Open Biosystems |
| KNY140 | *MAT***a** *ade2-1 ura3-1 his3-11,15 trp1-1 leu2-3,112 can1-100 pdr5Δ::HPH pep4Δ::LEU2* | (1) |
| KNY220 | *MAT***a** *ade2-1 ura3-1 his3-11,15 trp1-1 leu2-3,112 can1-100 pdr5Δ::HPH pep4Δ::LEU2 HRD1-3FLAG-* $KAN^R$ | (1) |

**Supplementary Table II**

Plasmids used in this study

| Name | Description | Source |
|---|---|---|
| pKN12-22 | CPY*-3HA, *CEN/ARS*, *URA3*<br>(CPY*: a soluble substrate due to the presence of a missense mutation in an otherwise vacuole-targeted protease) | (2) |
| pSM70 | KHN-3HA, *CEN/ARS*, *URA3*<br>(KHN: a heterologously expressed simian virus 5 hemagglutinin neuraminidase (HN) that is fused with the cleavable signal sequence from the yeast Kar2 (the ER luminal Hsp70)) | (3) |
| pSM101 | KWW-3HA, *CEN/ARS*, *URA3*<br>(KWW: a chimeric protein comprising KHN luminal domain/Wsc1 transmembrane domain/Wsc1 cytosolic domain) | (4) |
| pKN66 | Ste6*-3HA, *CEN/ARS*, *URA3*<br>(Ste6*: a C-terminal truncated version of the a-factor transporter Ste6) | (5) |
| pKN515 | 3HA-Pca1, *CEN/ARS*, *URA3*<br>(Pca1: a cadmium transporting P-type ATPase whose proteasome-dependent degradation is exclusively dependent on Doa10) | (6) |
| pKN541 | 3HA-Pdr5*, *CEN/ARS*, *URA3*<br>(Pdr5*: a 12 transmembrane protein that harbors misfolded lesions near these domains) | This study |
| pKN562 | 6myc-Hmg2, *CEN/ARS*, *URA3*<br>(Hmg2: the yeast HMG-CoA reductase isozyme) | This study |
| pJC104 | 4×UPRE (four copies of the unfolded protein response element)-*lacZ*, *2μ*, *URA3* | (7) |

A plasmid encoding *3HA-PDR5** (pKN541) was constructed as follows. The PCR reaction was performed using a plasmid pKN44 (8), which encodes *1HA-PDR5**, as a template and using primers OKN2234 and OKN2235. The resulting linear fragment was transformed into Mach1 competent cell (Thermo Fisher Scientific). The circularized plasmids were mini-prepped from the transformants and the DNA sequence was performed to verify that the single HA tag was replaced with triple HA tag.

A plasmid encoding 6myc-Hmg2 under the sequence of the *GPD* promoter (pKN562) was constructed as follows. The DNA fragment encoding the open reading frame of 6myc-Hmg2 was amplified by PCR from yeast strain expressing this protein (9) using primers OKN2273 and OKN2275. The resultant fragment was digested with *XbaI*/*XhoI* and inserted into the same sites of p416GPD (10).

**Supplementary Table III**

Oligonucleotide primers used in this study

| Name | Sequence |
|---|---|
| OKN2234 | CCTTATGATGTCCCAGATTACGCAGGTTCTTATCCTTACGA TGTACCAGACTACGCCGGTCCCGAGGCCAAGCTTAACAAT AACG |
| OKN2235 | CCTGCGTAATCTGGGACATCATAAGGGTAACCAGCATAAT CAGGAACGTCATAAGGGTATGAACCCATTTTTGTCTAAAGT CTTTCG |
| OKN2273 | ATATCCTCGAGCACCATGTAAACTACAAGAG |
| OKN2275 | GCTGCTCTAGAATGTCACTTCCCTTAAAAACGATAG |

**Supplementary Table IV**

Antibodies used in this study

| Antibody | Company | Identifier or reference |
|---|---|---|
| αHA | MEDICAL & BIOLOGICAL LABORATORIES | #M180-3 |
| αPgk1 | abcam | [22C5D8] (#ab113687) |
| αHrd1 | In house | (1) |
| αCdc48 | In house | (1) |
| αYos9 | In house | (1) |
| αDer1 | In house | (1) |
| αHrd3 | Gift from Dr. Thomas Sommer and Dr. Ernst Jarosch (Max-Delbrück-Center for Molecular Medicine, Berlin, Germany) | |
| αSic1 | Gift from Dr. Takumi Kamura (Nagoya University, Aichi, Japan) | |
| αClb2 | Gift from Dr. Takumi Kamura (Nagoya University, Aichi, Japan) | |
| αmyc | Gift from Dr. Takumi Kamura (Nagoya University, Aichi, Japan) | |
