## Supplemental Figure S1 for "Hydroxyurea inhibits ERAD-L independently of S-phase arrest in budding yeast"

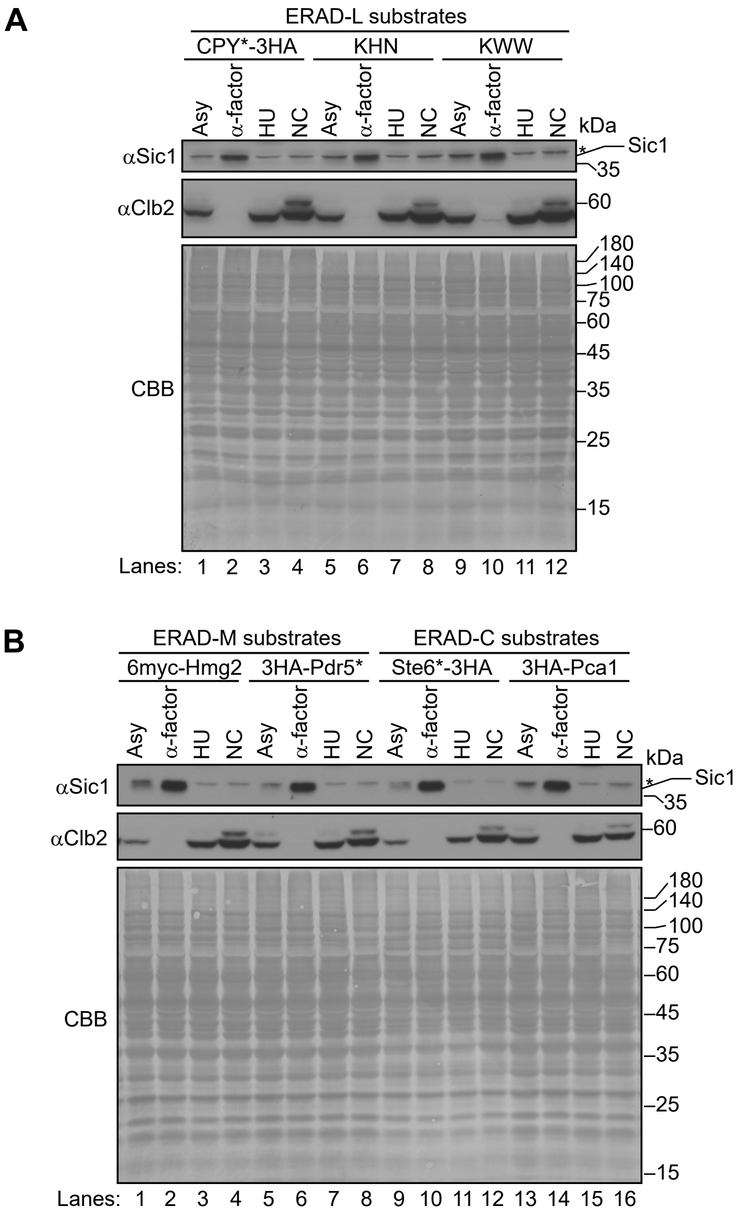


**Fig. S1: The levels of Sic1 and Clb2 in cells expressing ERAD-L, -M, or -C substrates synchronized at G1, S, or G2/M phase.**

(A-B) The levels of Sic1 and Clb2 in cells at time=0 of cycloheximide chase analysis in Fig. 1 and 2 were analyzed by western blotting. CBB staining of the membrane served as a loading control.

**Takano and Nakatsukasa, Figure S1**
