## Supplemental Figure S2 for "Hydroxyurea inhibits ERAD-L independently of S-phase arrest in budding yeast"

A

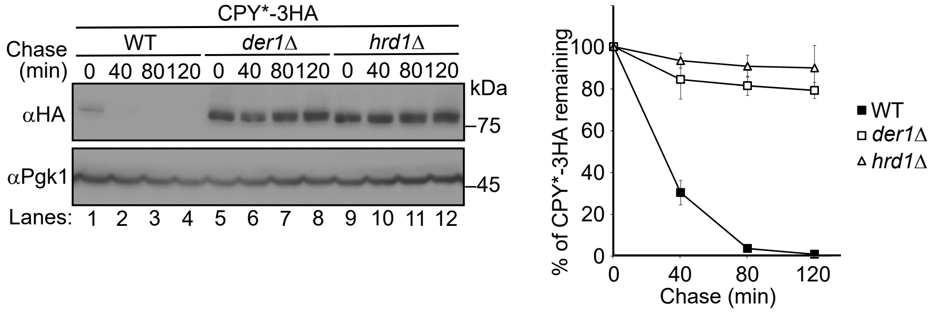


B

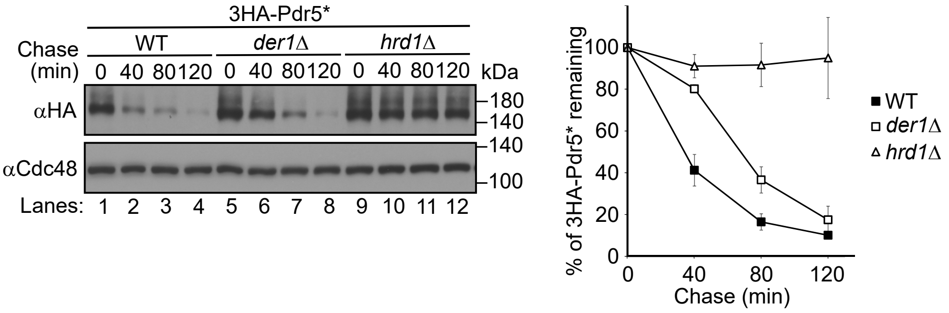


**Fig. S2: Der1 is dispensable for 3HA-Pdr5* degradation**

(A-B) Cycloheximide chase analyses of CPY*-3HA and 3HA-Pdr5* in WT, *der1*Δ, and *hrd1*Δ cells were performed as in Fig.1B. Signals for each substrate were normalized to Pgk1 signals (A) or Cdc48 signals (B), respectively. The data represent the mean ±SE of three independent experiments.

**Takano and Nakatsukasa, Figure S2**
