## Supplemental Figure S3 for "Hydroxyurea inhibits ERAD-L independently of S-phase arrest in budding yeast"

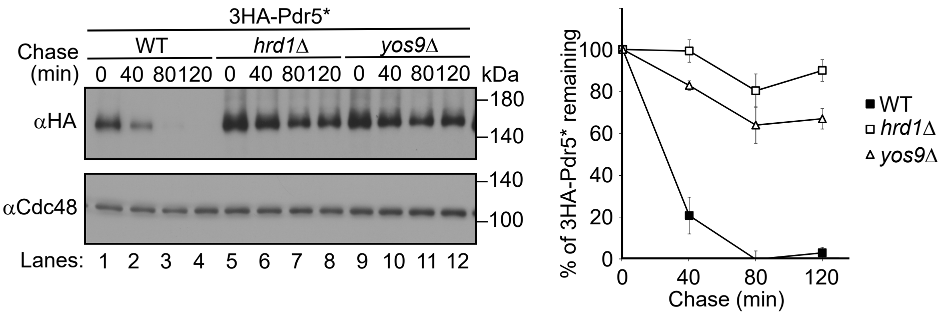


**Fig. S3: Yos9 is required for 3HA-Pdr5* degradation**

Cycloheximide chase analysis of 3HA-Pdr5* in WT, *hrd1*Δ, and *yos9*Δ cells was performed as in Fig.1B. Signals for each substrate were normalized to Cdc48 signals. The data represent the mean ±SE of three independent experiments.

**Takano and Nakatsukasa, Figure S3**
